# SpatialPrompt: spatially aware scalable and accurate tool for spot deconvolution and clustering in spatial transcriptomics

**DOI:** 10.1101/2023.09.07.556641

**Authors:** Asish Kumar Swain, Vrushali Pandit, Jyoti Sharma, Pankaj Yadav

## Abstract

Spatial transcriptomics has advanced our understanding of tissue biology by enabling sequencing while preserving spatial coordinates. In sequencing-based spatial technologies, each measured spot typically consists of multiple cells. Deconvolution algorithms are required to decipher the cell-type distribution at each spot. Existing spot deconvolution algorithms for spatial transcriptomics often neglect spatial coordinates and lack scalability as datasets get larger. We introduce SpatialPrompt, a spatially aware and scalable method for spot deconvolution as well as domain identification for spatial transcriptomics. Our method integrates gene expression, spatial location, and single-cell RNA sequencing (scRNA-seq) reference data to infer cell-type proportions of spatial spots accurately. At the core, SpatialPrompt uses non-negative ridge regression and an iterative approach inspired by graph neural network (GNN) to capture the local microenvironment information in the spatial data. Quantitative assessments on the human prefrontal cortex dataset demonstrated the superior performance of our tool for spot deconvolution and domain identification. Additionally, SpatialPrompt accurately decipher the spatial niches of the mouse cortex and the hippocampus regions that are generated from different protocols. Furthermore, consistent spot deconvolution prediction from multiple references on the mouse kidney spatial dataset showed the impressive robustness of the tool. In response to this, SpatialPromptDB database is developed to provide compatible scRNA-seq references with cell-type annotations for seamless integration. In terms of scalability, SpatialPrompt is the only method performing spot deconvolution and clustering in less than 2 minutes for large spatial datasets with 50,000 spots. SpatialPrompt tool along with the SpatialPromptDB database are publicly available as open source <u>software</u> for large-scale spatial transcriptomics analysis.

## Introduction

Recent advances in sequencing technologies enable researchers to sequence and analyse samples up to the single-cell resolution. Although scRNA-seq technologies could reveal the heterogeneity of cell-types and cell states, but they fail to provide any spatial information^1^. In recent years, several spatially resolved transcriptomics approaches have emerged that preserve the spatial information of cells while measuring the gene expression profiles^2–5^. Spatial transcriptomics methods can be broadly classified into two categories, namely, fluorescence in situ hybridisation (FISH)-based and sequencing-based^6^. FISH-based methods allow high resolution spatial localisation of transcripts but have limited ability to detect large numbers of genes^7,8^. Conversely, sequencing-based methods, such as 10X Visium^3^ and ST^5^ can detect thousands of genes but have low resolution (10-100 μm). In low-resolution spatial techniques, each measured location, referred as spot, usually contains mixture of cell-types^8^. Recently, high-resolution sequencing-based techniques such as Slide-seq^4^ and HDST^9^ measure each spot typically consists of 1-2 cells. However, the fixed beads applied in these techniques cannot perfectly match the boundaries of individual cells, thus resulting in multiple cells being captured within a single spot^10^. Each spot in the sequencing-based spatial methods usually contains a mixture of cell-types. Therefore, identifying the cell-type distribution at each spot is a major challenge in understanding the complex tissue architectures.

Recent spot deconvolution tools have leveraged the cell-type specific gene signatures from reference scRNA-seq datasets to perform spot deconvolution in spatial datasets. For instance, SPOTlight^11^ and CARD^12^ tools use a non-negative matrix factorisation approach to deconvolute the spots based on cell-type specific marker genes from the scRNA-seq reference dataset. In contrast, the RCTD^13^ tool uses a supervised approach that employs a probabilistic model to derive maximum likelihood estimate of the proportions of different cell-types. Cell2location^14^ and Stereoscope^15^ tools rely on the assumption that both spatial and scRNA-seq datasets follow a negative binomial distribution. They utilise this assumption as a foundation to predict the cell-type inference in the spatial data. Recently, the Tangram^16^ tool adopts a deep learning framework to align the scRNA-seq data as ‘puzzle pieces’ to match the spatial gene expression patterns measured by spatial technologies while correcting for inter-platform differences. None of these available tools, except CARD, have exploited the rich spatial information available in the spatial datasets. Most of these ignore the impact of the microenvironment and important biological factors useful in the context of tissue arrangement. In a typical spatial tissue arrangement, nearby spots tend to have similar cell-type compositions compared to the distant spots. Moreover, the cellular microenvironments play a major role in determining the cell-type composition of spatial spots^17,18^. To achieve realistic cell-type inference, it is necessary to perform spatially informed spot deconvolution of the spatial datasets while considering the effects of microenvironment and spatial arrangement.

Furthermore, the identification of spatial domains that accurately reflect biological reality remains a challenging task in spatial transcriptomics. Most domain identification methods can be grouped into two categories, namely, non-spatial and spatial-based methods^19^. The non-spatial-based methods donot utilise the spatial coordinate information of the spatial datasets. Most of them are merely an extension of the tools that are commonly used for scRNA-seq datasets (e.g., Seurat^20^ and Scanpy^21^). Recent spatial-based methods such as BayesSpace^22^, SpaGCN^23^, STAGATE^24^, and SEDR^25^ can predict spatial domains based on spatial locations and tissue architecture. While spatial-based methods are biologically interpretable in identifying spatial domains compared to their non-spatial counterpart, many of them have scalability issue for large spatial datasets. In addition, spatial-based methods rely on computationally intensive algorithms such as graph convolution networks and variational graph auto-encoder, which becomes challenging to apply on large datasets^26,27^. Moreover, the spatial-based clustering methods neglect the potential influence of the local microenvironment on gene expression. Many of these methods assume that cells within a given spatial domain are positioned close together^28^. This assumption, however, ignores the fact that distant cells can be part of the same microenvironment^29,30^ and nearby cells (i.e., rare cell groups) can also be part of two different microenvironments.

Here, we introduce a spatially aware scalable and accurate tool, referred as SpatialPrompt, for spot deconvolution and clustering in spatial transcriptomics. SpatialPrompt utilises a state-of-the-art spatial simulator to generate spatial spots that closely resemble the real spatial data. SpatialPrompt uses an iterative approach inspired from the message passing layer in the graph neural net (GNN)^31^ to compute weighted mean expression of the local microenvironment for each spatial spot. By encoding the spot’s own expression and local weighted mean expression, a concatenated matrix is created that can be used for domain identification in spatial data. An integrated model was constructed for spot deconvolution that combines non-negative ridge regression (NRR) with K-nearest neighbour (KNN) regressor model. This model will deconvolute the real spatial spots by incorporating spatial embeddings into the simulated spatial spots. SpatialPrompt is highly scalable and able to perform spot deconvolution and clustering on a large dataset containing 50,000 spatial spots within 120 seconds. We demonstrated the sensitivity and accuracy of SpatialPrompt for spot deconvolution and domain prediction using mouse cortex and hippocampus datasets generated from 10X Visium^3^ and Slide-seq^4^ platforms. Furthermore, a quantitative assessment is performed using the gold standard annotations from the human dorsal prefrontal cortex (DLPFC) spatial dataset^32^. To further test the robustness of the SpatialPrompt, we perform spot deconvolution on the mouse kidney spatial dataset using two publicly available unpaired scRNA-seq reference datasets^33,34^. Additionally, we built an open-access scRNA-seq database having 41 manually annotated reference datasets to facilitate seamless integration during spatial deconvolution.

## Materials and Methods

### Datasets and pre-processing

We retrieved four publicly available spatial datasets and their corresponding scRNA-seq reference datasets from multiple databases. All the spatial and scRNA-seq datasets were pre-processed using the *Scanpy* package^21^. Low-quality cells (or spots) and genes were removed from both spatial and scRNA-seq datasets. The details of the four datasets used in this work are described below.

#### Human prefrontal cortex dataset

This spatial dataset was retrieved from the Lieber Institute for Brain Development (LIBD) database^32^. This dataset has been generated through the sequencing of 12 tissue slides obtained from the human DLPFC region. Following the previous studies^19,35,36^, the tissue slide numbered 151673 was considered for the performance assessment of our tool. This chosen slide comprised 3,639 spots and 33,538 genes. This spatial dataset has gold standard manual annotations of seven cortical layers (layer-1 to layer-6 and white matter), which is used as a ground truth for comparative evaluation of our method. The corresponding reference single-nucleus RNA sequencing (snRNA-seq) dataset comprising 78,886 cells and 30,062 genes was retrieved from the GEO database (accession id: GSE144136^37^).

#### Mouse 10X Visium cortex dataset

This spatial dataset was collected from the 10X Visium database, which was sequenced from the mouse cortex^3^. This dataset has 2,559 spots and 1,337 genes for the anterior section. The scRNA-seq dataset of the moue cortex dataset was obtained from GEO database having accession id: GSE71585^38^. The scRNA-seq dataset has 14249 cells and 34617 genes. In the Visium spatial technique, mRNAs were captured from the beads having diameter of 55 μm and bead-to-bead distance of 100 μm^3^. These low-resolution spots contain multiple cells (e.g., 10 to 50) depending on the cell-type and cell-class (e.g., normal or tumour).

#### Mouse hippocampus slide-seq dataset

This spatial dataset was obtained from the mouse hippocampus, which has been sequenced using the Slide-seq technique^4^. This large spatial dataset has 53,173 spots and 23,264 genes. The Slide-seq spatial dataset, along with the corresponding reference scRNA-seq data, were retrieved from the single cell portal^13^ hosted by the Broad Institute. The sc-RNAseq data comprises 52846 cells and 27953 genes. Slide-seq is a high-resolution technique using a bead diameter of 10 μm, and each spot comprises of 1-2 cells^4^.

#### Mouse Visium kidney dataset

This spatial dataset was retrieved from mouse kidney and was available at the STOmicsDB^39^ database. In this dataset, all five slides were sequenced in a five-time frame. This dataset has 1,617 spots and 32,285 genes. In addition, two mouse kidney scRNA-seq datasets were retrieved from GEO having accession numbers GSE157079 and GSE107585^33,34^. The GSE157079 dataset has 43,636 cells and 31,053 genes, and the GSE107585 dataset has 43,745 cells and 16,272 genes. The former dataset (GSE157079) was sequenced using the Illumina HiSeq 4000/3000 platform in the USA, while the latter was sequenced using the Illumina HiSeq 2000 platform in South Korea.

### Spatial spot simulation using reference scRNA-seq dataset

We simulated the spatial data from the reference scRNA-seq data using our own spatial simulator. This is because most of the existing spatial simulators simulate the spots by randomly merging the single cells^36^. In addition, these simulators often ignore the spots residing in the core of the microenvironment, which consist of one or two predominant cell types. Our spatial simulator uses a three-step process to simulate the spots. In step 1, the simulator generates spots that majorly comprise of one cell type, which resemble the spots that reside at the core of microenvironment. In step 2, it simulates spots that predominantly consist of 2-3 cell types, which resemble the spots located at the border of two microenvironments. Lastly, in step 3, the simulator randomly merges single cells to simulate spots that resemble the spots in the border of multiple microenvironments and rare cell types.

### Spatial and scRNA-seq data integration for spot deconvolution and clustering

The following notations will be used to describe the spot deconvolution and clustering model used in our SpatialPrompt tool:

*M*_*SC*_ : scRNA-seq matrix having *N*_*SC*_ cells as rows and G genes as columns,

*C*_*SC*_ : cell type annotations of the matrix *M*_*SC*_,

*M*_*SP*_ : spatial RNA-seq matrix having *N*_*SP*_ spots as rows and G genes as columns,

*x, y*: x and y coordinates of the matrix *M*_*SP*_,

*G*_*C*_ : number of common genes between *M*_*SC*_ *and M*_*SP*_,

*G*_*H*_ : top high variance common genes identified from the matrix *M*_*SC*_,

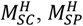 : single cell and spatial matrices after indexing *G*_*H*_ genes from the matrix *M*_*SC*_ and *M*_*SP*_,

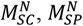: Counts per million normalised single cell and spatial matrices,

*M*_*SI*_ : simulated spatial matrix having *N*_*SI*_ number of spots as rows and M_H_ genes as columns,

*C*_*SI*_: known cell-type proportion matrix of the simulated spot matrix *M*_*SI*_,

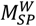 local weighted mean expression of all spatial spots from the matrix *M*_*SI*_,

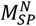 column-wise concatenated matrix of 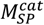 and 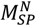 having dimension of *N*_*SP*_*× 2G*_*H*_,

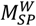: predicted local weighted mean expression of matrix *M*_*SI*_,

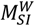: concatenated matrix of *M*_*SI*_ and 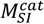 having dimension of *N*_*SI*_*× 2G*_*H*_.

Our SpatialPrompt tool requires *M*_*SC*_ and *M*_*SP*_ matrix along with *x, y* coordinate information and cell type annotation *C*_*SC*_ for the spot deconvolution. On the other hand, for domain identification, SpatialPrompt need only the matrix *M*_*SP*_ along with the coordinate information. At first, our tool identifies the *G*_*C*_ common genes between matrix *M*_*SC*_ and *M*_*SP*_. Of these *G*_*C*_ common genes, the *G*_*H*_ (default value is 1,000) number of high-variance genes is extracted from matrix *M*_*SC*_. After this, matrices 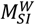 and 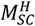 are created by indexing *G*_*H*_ genes from the matrix *M*_*SC*_ and *M*_*SP*_. Next, counts per million normalisation was applied to 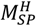 and 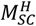 as follows:

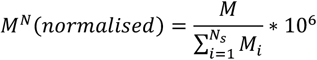

Here, *M*^*N*^ represent the normalised matrix of M having N_S_ cells or spots. Next, the 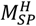 was used to simulate the *M*_*SI*_ matrix with known cell type mixture (*C*_*SI*_) having *N*_*SI*_ spots using the three-tier steps as described above. By default, *N*_*SI*_ was set within the range of 20,000 to 30,000. To set the optimum value of *N*_*SI*_, benchmarking analysis was performed by simulating spatial spots in the range from 10^3^ to 10^6^ using the human DLPFC spatial dataset. We observed that SpatialPrompt performed consistently after *N*_*SI*_ reached 20,000. Therefore, the default value for *N*_*SI*_ was set between 20,000 and 30,000.

To incorporate the microenvironment information into the spatial matrix 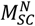, the weighted mean expression was calculated for each spatial spot using a three-step process (see **Fig. 1B**). In step 1, K nearest neighbours were identified for each spot using the Scipy *cKDTree* spatial module^40^. The *cKDTree* module was used as it identifies neighbours very fast by dividing the search space into smaller regions with hyperplanes and searching only the spots that are likely to contain the nearest neighbour^40,41^. Moreover, it avoids calculating distances to many points that are relatively far away from the query point, thus making its application ideal for larger datasets as in our case. In step 2, for each spot (*S*_*Q*_), the cosine similarity was calculated with its K neighbours (*S*_*K*_) where, *S*_*K*_ : {*S*_1_, *S*_2,_… *S*_*K*_} follows:

**Fig. 1:**
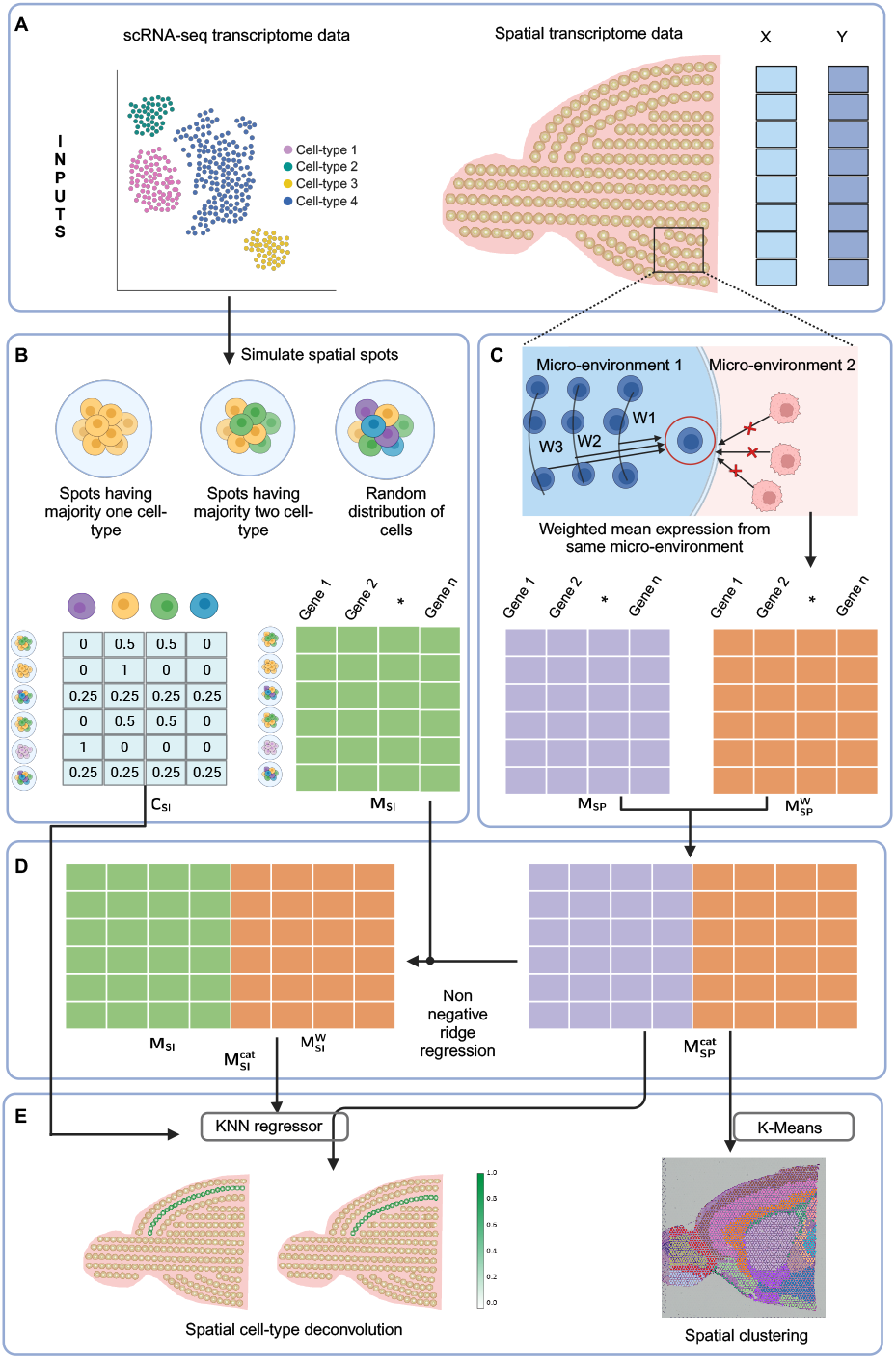
Overall workflow of SpatialPrompt. (A) SpatialPrompt takes spatial matrix with coordinate information and scRNA-seq matrix with cell type annotations as input for spot deconvolution and clustering. (B) The spatial spot simulation pipeline utilises scRNA-seq expression matrix and cell type annotations to generate simulated expression matrix (*M*_*SI*_) with known cell type mixture (*C*_*SI*_). Three criteria were used to generate the *M*_*SI*_ . (C) Using X and Y coordinates of spatial matrix (*M*_*SP*_), weighted mean expression 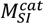 was calculated for each spatial spot in the real spatial data. (D) The non-negative ridge regression model is employed that predicts the weighted mean neighbour expression 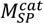 of the simulated spatial matrix (*M*_*SI*_) by utilising real spatial expression (*M*_*SP*_) and its weighted mean neighbour expression 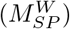. (E) SpatialPrompt outputs spatial clustering using K-means on concatenated spatial matrix 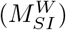. For spatial deconvolution KNN regressor model trained on 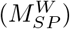 and (*C*_*SI*_) and predicts cell type proportions on real spatial matrix 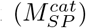.

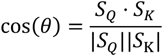

Here, cos(θ) represents the cosine similarity, *S*_*Q*_ *⋅ S*_*K*_ denotes the dot product of vectors, |*S*_*Q*_| and |*S*_K_| represents the magnitude (norm). After calculating the cosine similarities, only the neighbours having high similarity were considered for weighted mean expression calculation. To select highly similar neighbour spots, out of K neighbours, K/2 number of neighbours having higher cosine similarity were selected for further steps. In step 3, an iterative approach inspired from the message-passing layer in the graph neural net (GNN) was applied to calculate the weighted mean expression matrix 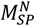 for each spot as follows:

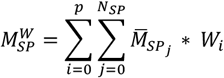

Here, 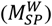is the mean expression of the neighbours of the j^th^ spatial spot, *p* is the number of iterations, message passing layer will incorporate weighted mean into the spot. *W*_*i*_ is the weight assigned to the mean expression in the i^th^ iteration. In each iteration, *W*_*i*_ is reduce by *W*_1_/i (i.e., *W*_*i*_ = 1 at i = 1) so that higher weightage will be assigned to the nearer spots as compared to farther spots. It is recommended to set the value of parameter *p* between 5 to 10, as after 5th iterations all the spots will have the weighted mean expression of their neighbours from the same microenvironment. Next, matrices 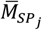 and 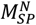were concatenated column-wise to create the matrix 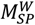. The matrix 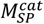 was further used for domain identification by applying k-means clustering algorithm.

### Non-negative ridge regression (NRR) model

As the simulated spatial data (*M*_*SI*_) does not have spatial coordinate information, a NRR model was built to predict the local weighted mean expression. This model predicts the matrix 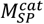 based on 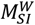 and 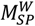. The regression coefficient in the ridge regression model was obtained by minimising the cost function:

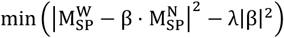

Here, λ is the penalty or alpha parameter and β denotes the regression coefficient of the ridge regression model. To determine the optimum λ, a subset of 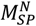and 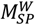was selected for the hyperparameter tuning. After obtaining the β on the optimised λ, *M*^’^ can be predict by,

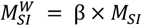

In the next step, *M*_*SI*_ and 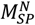were concatenated to form the matrix 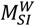. As 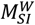have information of own expression and predicted weighted mean expression of all simulated spots with known cell type proportions *C*_*SI*_. A KNN-regressor model was trained using the matrix 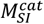and *C*_*SI*_ as target. Here for each real spatial spot in 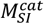, the KNN-regressor model^42^ will find the K most related simulated spots in the kernel space with least Minkowski distance and predicts the cell type proportions.

### Benchmarking Performance of SpatialPrompt

The gold standard annotations in the DLPFC dataset were used to assess the performance of SpatialPrompt with existing methods for both spot deconvolution and domain identification. For spot deconvolution, we compared our method with 5 different methods, namely, CARD, Cell2location, Tangram, SPOTlight and RCTD. Among these methods, CARD, Cell2location, and Tangram have been reported as the best-performing methods in a recent benchmarking study^49^. Further, SPOTlight and RCTD were the most cited methods for spot deconvolution. The 10 layer-specific excitatory neuron cell-types were retrieved from the reference sn-RNAseq retrieved from human brain cortex region. The area under receiver operating characteristics (AUROC) score was computed to assess the overall performance of all the methods. The AUROC analysis involved determining the sensitivity and specificity for classifying a spot belonging to a particular cell-type.

For benchmarking the domain identification performance of SpatialPrompt, we selected 6 different tools *viz*. BayesSpace, SEDR, STAGATE, SPAGCN, Scanpy, and Seurat. The accuracy of domain identification tools was evaluated by calculating the normalised mutual information (NMI) score. This score measures the effectiveness of a tool by calculating the mutual information between the predicted clusters and the gold standard annotations. The NMI score was calculated between predicted clusters (*Y*_*P*_) and ground truth annotations (*Y*_*R*_) using,

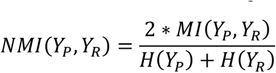

Here, *MI*(*Y*_*P*_, *Y*_*R*_) in the numerator denotes the mutual information between *Y*_*P*_ and *Y*_*R*_, while *H*(*Y*_*P*_), *H*(*Y*_*R*_) represent the entropy of the *Y*_*P*_ and *Y*_*R*_, respectively.

The time complexity was calculated using the hippocampus slide-seq dataset. This dataset was also chosen to assess the scalability of all the methods, because it consists of 53,173 spatial spots. Four spatial datasets were randomly sliced from the slide-seq dataset containing 5,000, 10,000, 20,000, and 50,000 spots. Since most of the previous spatial deconvolution tools were not designed for large reference datasets, the scRNA-seq reference dataset was downsampled to 13,247 cells and 10,000 genes. For down-sampling the reference dataset, the cell-types having counts higher than 1,000 were trimmed down to 1,000 by randomly selecting cells and 10,000 high-variance genes were selected for the downstream analysis. Additionally, for all the tools, if the option to select the number of high-variance gene was available, it was set between 1,000 to 2,000 or set to the number recommended by the method. The following parameters were used for benchmarking as described below for each method:

We followed the instructions available in the documentation of respective tools and used the default parameter unless mentioned explicitly. For *Cell2location*, due to the heavy computational burden, for 20,000 and 50,000 spots, max_epochs parameter was reduced to 2,000 (default value: 30,000) while other parameters were set at default values. Clustering pipeline provided by STAGATE, SEDR, and SpaGCN on their GitHub repository was followed using default parameters. For Scanpy, all the datasets were loaded into the tool followed by application of counts per million and log-normalisation to the data. To obtain clusters in the Scanpy, the command *pp*.*neighbors* was used. For Seurat, SCTransform normalisation was applied to the spatial data after basic quality control. To obtain the clusters, the *FindClusters* command was used with a resolution of 0.5. The analysis was performed on a system with an Intel Xeon processor with 48 cores, 128 GB of RAM, and 4 GB of graphics memory.

### SpatialPromptDB database

We created a SpatialPromtDB database using the Mkdocs client^43^ to store the scRNA-seq reference datasets along with their cell-type annotations. The scRNA-seq datasets of major tissue classes were obtained from different databases, including GEO^44^, human cell atlas^45^, single cell portal^46^ of Broad Institute and Figshare^47,48^. At first, all these downloaded scRNA-seq datasets were loaded and pre-processed using the *Scanpy* package. Cells with less than 500 reads and genes having total reads of less than 1000 were removed. Additionally, cells having high mitochondrial genes (above 20%) were excluded. Next, counts per million normalisation and logarithmic transformation were applied to the datasets. If the cell annotations were not provided by the original study, the clusters were obtained using the *pp*.*neighbors* function available in the *Scanpy*. Cell type or cluster-specific markers were identified using the *tl*.*rank_genes_groups* function. These markers were further validated manually using the CellMarker^49^ and PanglaoDB^50^ databases. Finally, the raw count matrix of the expression data and cell-type annotations were exported in tabular-form from the *Scanpy*. Finally, all the reference datasets with their cell-type annotations were hosted on SpatialPromptDB database (https://swainasish.github.io/SpatialPrompt).

## Results

### Overview of SpatialPrompt

SpatialPrompt framework learns the cell-type specific gene signatures from the scRNA-seq reference dataset and performs spatially informed spot deconvolution. The framework of SpatialPrompt tool operates in 4-tier manner. At first, a spatial simulator simulates spatial spots from the reference scRNA-seq dataset using various criteria to mimic the real spatial data (**Fig. 1B**). The basic idea behind the spatial simulator is that spots residing in the core of the microenvironment predominantly consist of one cell type and spots reside at the border of microenvironments have mixture of cell types. In the second step, an iterative approach inspired by the message-passing layer in the GNN was applied to incorporate the local microenvironment information into the spatial data. This step calculates the weighted mean expression of the local microenvironment for each spatial spot. Next, a concatenated representation of the spatial data was created by combining the spot’s own expression and local weighted mean expression (**Fig. 1C**). Thus, this concatenated expression matrix captures the spot’s individual characteristics along with the local microenvironment information. In the third step, a NRR model was fit to concatenated matrix to learn the local microenvironment gene expression patterns from the real spatial data. This model was employed to predict the spatial pattern in the simulated spatial data (**Fig. 1D**). In the last step, k-means clustering is applied on the concatenate matrix for domain identification. Moreover, for spot deconvolution, a K-nearest neighbour (KNN) regressor model is utilised (**Fig. 1E**). This model learns the cell type specific gene signatures from the spatial embedded simulated matrix and predicts the cell type proportions in the real spatial data.

### Benchmarking Performance of SpatialPrompt

The performance of SpatialPrompt was evaluated using the human DLPFC spatial dataset. At first, both real spatial data and their corresponding reference scRNA-seq datasets were integrated. Then, 5 different methods were applied on this integrated dataset to calculate the cell-type proportions in the spatial spots. Performance of SpatialPrompt for spot deconvolution compared with five other methods,: namely, CARD^12^, Cell2location^14^, Tangram^16^, SPOTlight^11^, and RCTD^13^. The layer discriminative accuracy of the predicted cell proportion for all methods was assessed by calculating the AUROC score using layer annotation as the ground truth. Our analysis shows that SpatialPrompt was able to capture the cell type topography for all 10 excitatory neurons (**Fig. 2A & 2B)**.

**Fig. 2:**
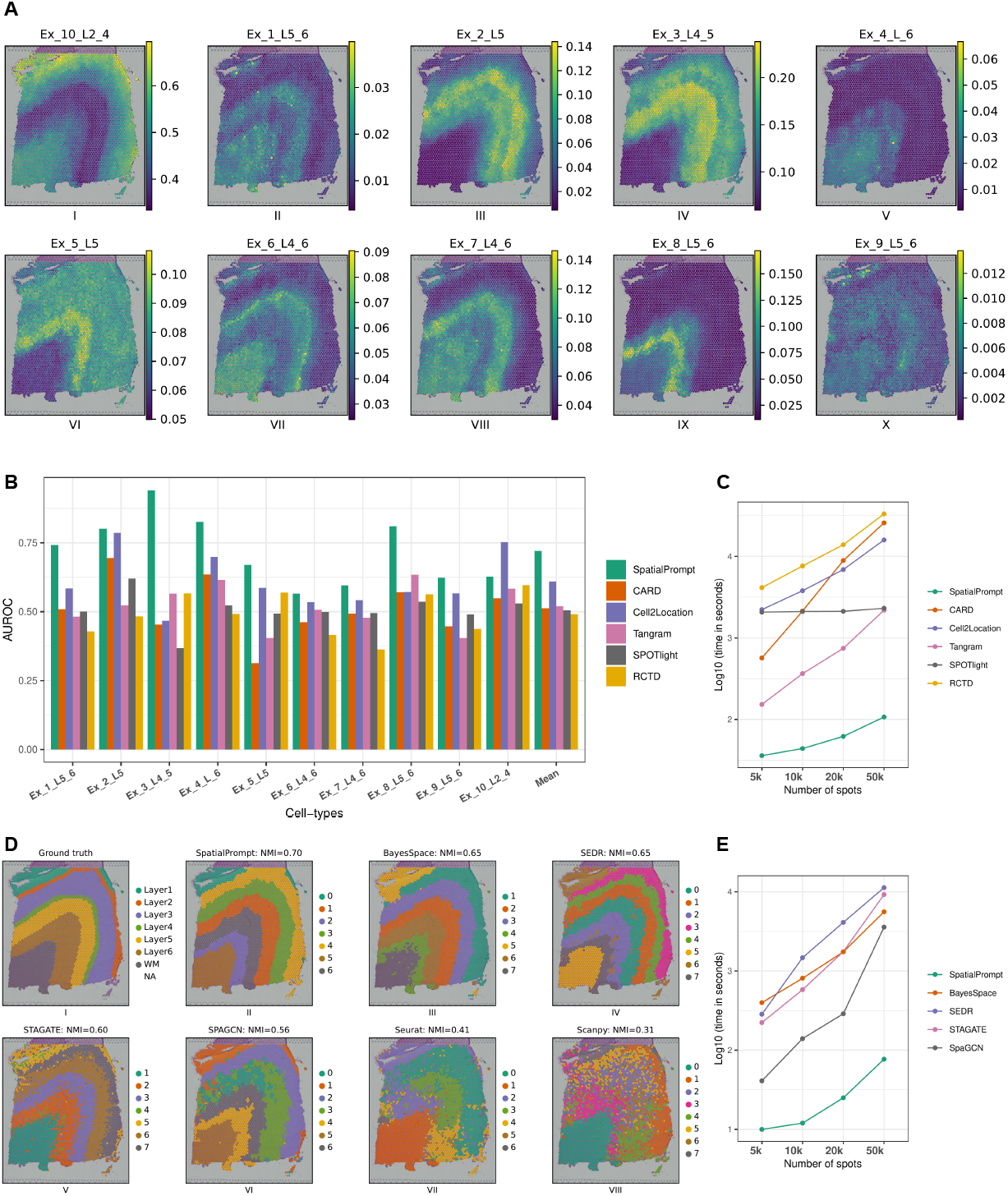
Quantitative assessments of spot deconvolution and clustering on various tools on the human DLPFC dataset. (A) Cell-type distribution of 10 major neuronal cell types predicted by SpatialPrompt on DLPFC dataset. (B) Bar plot of AUROC scores obtained by comparing gold standard annotations of seven layers (Fig. 2D(I)) with five other deconvolution tools. Higher AUROC scores indicate better accuracy and performance. Running time by six deconvolution tools to process 5,000 to 50,000 spatial spots. The x-axis represents the number of spatial spots, and the y-axis represents the logarithmic scale of the running time for each tool. Clustering performance of SpatialPrompt and six other tools (III-VIII). The NMI score is used to measure the similarity between the ground truth (I) and the clusters predicted by each method. Higher NMI scores indicate more accurate clustering. (E) Running time by SpatialPrompt and four other spatial-based clustering tools to process 5,000 to 50,000 spatial spots. The x-axis represents the number of spatial spots, and the y-axis represents the logarithmic scale of the running time for each tool.

SpatialPrompt showed superior performance compared to all existing methods with a mean AUROC score of 0.72 (**Fig. 2B**). Cell2location was the second-best method next to SpatialPrompt with a mean AUROC score of 0.60. The AUROC scores for remaining 4 methods were as follows: Tangram (0.52), CARD (0.51), Spotlight (0.50), and RCTD (0.49) (**Fig. S3-6**).

The total running time of all methods was calculated using the hippocampus slide-seq dataset, which comprising 53,173 spots and 23,264 genes. This spatial dataset was used to create four low-resolution spatial subsets with varying number of spots, as described in the method section. Again, SpatialPrompt exhibited superior performance with up to 20 times faster as compared to all other benchmarked methods (**Fig. 2C & Table. 3**). Noteworthy, SpatialPrompt showed high scalability by efficiently handling thousands of spatial spots. Indeed, SpatialPrompt is the only method capable of performing spatial deconvolution on a dataset comprising 50,000 spots in under 2 minutes. Except Tangram tool, all the 4 methods exhibited runtime of over 35 minutes to handle even a smaller spatial dataset containing only 10,000 spatial spots. The RCTD tool performed the worst among compared methods with runtime longer than 2 hours (**Table. 3**).

**Table 1.**
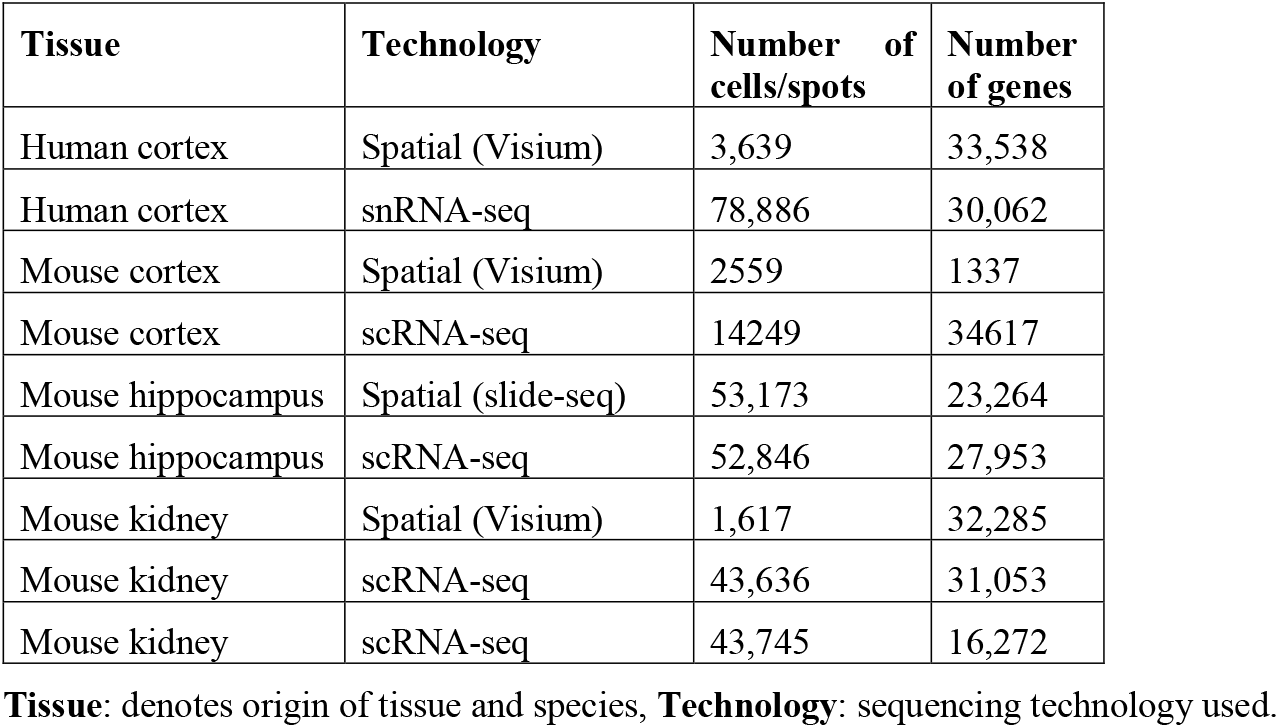
Summary of the datasets used in this study.

**Table 2.** Summary of the tools used in the benchmarking study.

| Tools | PMID | Method type |
| --- | --- | --- |
| **CARD** | 35501392 | Spot deconvolution |
| **Cell2location** | 35027729 | Spot deconvolution |
| **Tangram** | 34711971 | Spot deconvolution |
| **SPOTlight** | 33544846 | Spot deconvolution |
| **RCTD** | 33603203 | Spot deconvolution |
| **STAGATE** | 35365632 | Domain identification |
| **SEDR** | - | Domain identification |
| **SpaGCN** | 34711970 | Domain identification |
| **BayesSpace** | 34083791 | Domain identification |
| **Seurat** | 29608179 | Domain identification |
| **Scanpy** | 29409532 | Domain identification |

**Tools:** Name of tools used for benchmarking spot deconvolution and domain identification performance, **PMID:** PubMed IDs of tools, the notation ‘-’ refers to the tools unpublished or not available on PubMed.

**Table 3.** Summary of quantitative assessments obtained for spot deconvolution by comparing SpatialPrompt to five other tools.

| Tools | AUROC | T1 | T2 | T3 | T4 |
| --- | --- | --- | --- | --- | --- |
|  |  | (Ns:<br>5,000) | (Ns:<br>10,000) | (Ns:<br>20,000) | (Ns:<br>50,000) |
| **SpatialPrompt** | 0.72 | 36 | 44 | 62 | 107 |
| **CARD** | 0.51 | 568 | 2139 | 8892 | 25797 |
| **Cell2location** | 0.61 | 2225 | 3793 | 6894 | 15915 |
| **Tangram** | 0.52 | 153 | 365 | 745 | 2189 |
| **SPOTlight** | 0.51 | 2073 | 2109 | 2119 | 2310 |
| **RCTD** | 0.49 | 4140 | 7632 | 13896 | 33192 |

**Tools:** Name of tools used for benchmarking deconvolution performance, **AUROC:** Area under the receiver operating characteristics scores obtained by comparing the tool's output to the ground truth, **T1-T4:** Running time (in seconds) by all tools to process 5,000 (T1) to 50,000 (T4) spatial spots, **Ns:** represent the number of spatial spots.

**Table 4.**
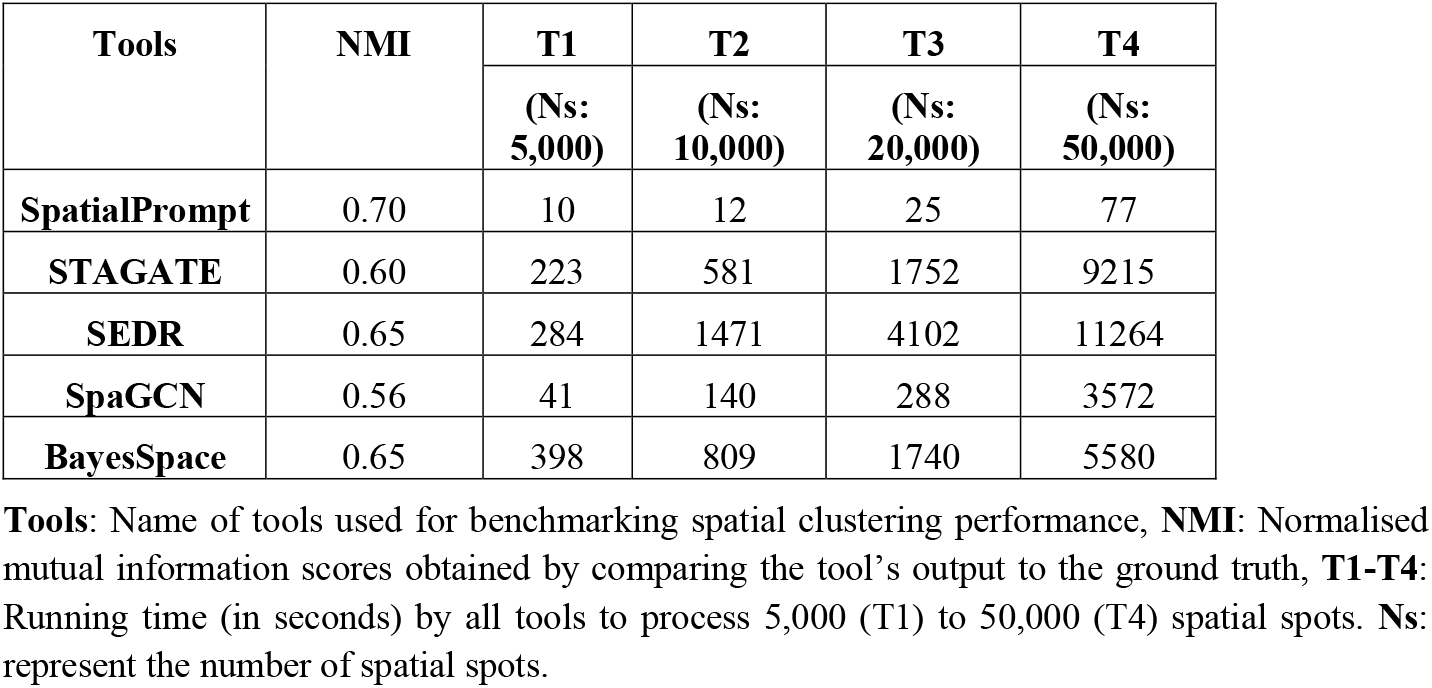
Shows the quantitative assessment result for domain identification.

The domain identification performance was assessed using slide 151673 of the DLPFC dataset. To benchmark the performance of our tool, we selected 4 recent spatial-based and 2 non-spatial based clustering techniques. The spatial-based tools included in this study were BayesSpace^22^, SEDR^25^, STAGATE^24^ and SPAGCN^23^ while Seurat^20,52^ and Scanpy^21^ were included as the non-spatial based tools. Interestingly, SpatialPrompt achieved the highest NMI score of 0.70, indicating its superior performance compared to the others (**Fig. 2D**). Among the spatial-based tools, only BayesSpace and SEDR performed well with an equal NMI score of 0.65. The remaining 2 methods had NMI score ≤ 0.60. In contrast, non-spatial methods *viz. Seurat* and *Scanpy* performed poorly with NMI scores 0.41 and 0.31, respectively. Furthermore, all the 4 spatial-based methods showed poor scalability as the number of spots increased greater than 5,000 (**Fig. 2E**). In contrast, SpatialPrompt could successfully identify domains in spatial dataset with 50,000 spots within 90 seconds (i.e., 44 to 150 times faster).

### Application of SpatialPrompt on mouse brain Visium and slide-seq datasets

The performance and scalability of the SpatialPrompt was additionally assessed using the mouse cortex Visium and hippocampus slide-seq datasets. These datasets were pre-processed as described in the method section. We applied SpatialPrompt and Cell2location tools on both datasets separately for spot deconvolution. Here, Cell2location was chosen as it was the second-best performing method after SpatialPrompt in our benchmark study.

The excitatory neurons in the mouse cortex are arranged sequentially in six layers labelled as L1 to L6 (**Fig. 3A**). Each layer has specific gene signatures and molecular function^53^. The layers L1 to L2/3 are rich in synaptic connections that facilitate intracortical communication^54^. The layer L4 is the main receiver of inputs coming from the thalamus^55^. The layers L5 and L6 are the cortex deep layers that have major role in long-distance projections to thalamus, striatum, and spinal cord^56^. Noteworthy, SpatialPrompt was able to predict each cell type proportions in the Visium dataset more clearly as compared to Cell2location (**Fig. 3B** and **Fig. S1**). In addition, a clear sequential organisation of the major cell-types was observed (**Fig. 3B**). A recent study reported that most deconvolution tools applied to this Visium dataset falsely predict the L2/3 cell type in the centre of the cortex region (caudoputamen and nucleus accumbens area)^57^. We noticed the same incorrect prediction made by the Cell2location tool, as shown in **Fig. S1**. In contrast, SpatialPrompt predicted almost negligible presence of L2/3 cells in the cortical centre region (**Fig. 3B**).

**Fig. 3:**
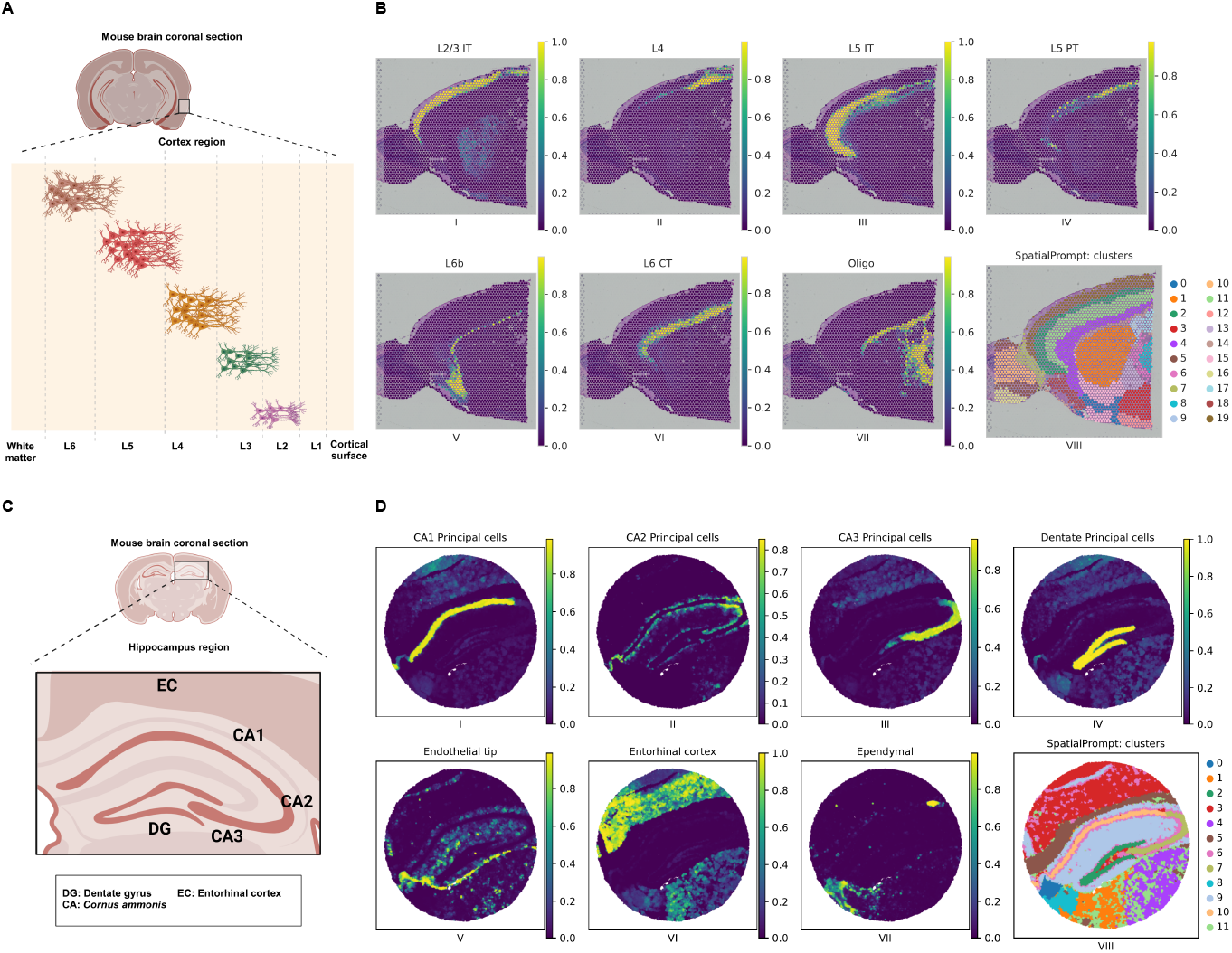
Spatial mapping of major cell types on mouse cortex and hippocampus. (A) Graphic illustration of sequential organisation of major cell types in mouse cortex from L1-L6 and white matter. (B) Spatial mapping of seven major cell types on mouse Visium cortex by SpatialPrompt. The figure on the bottom right (VIII) shows the clusters obtained by the SpatialPrompt (C) Application of SpatialPrompt on the mouse hippocampus Slide-seq dataset. The figure on the bottom right (VIII) shows the domains obtained by our tool.

Hippocampus majorly comprises of three regions: cornu ammonis 1-2 (CA1-2), cornu ammonis 3 (CA3) and denate gyrus (DG; **Fig. 3C**)^12,58^. The CA1 region is positioned closest to the subiculum, while CA3 is positioned adjacent to the DG. The CA2 is situated between CA1 and CA3. The DG has a “denate” or tooth-like appearance near CA3^59^. Entorhinal cortex (EC) lies in the medial temporal lobe, which is the main interface between the hippocampus and neocortex^60^. We noticed that SpatialPrompt precisely predicts the cell-types of all anatomical features especially for CA1, CA2, CA3, DG, and EC (**Fig. 3D**). In contrast, Cell2location was able to capture only a few of the anatomical features of certain cell types, such as CA3 and ependymal cells (**Fig. S2**). Despite runtime of 4.4 hours, the Cell2location tool incorrectly predicted some proportion of CA1 cells in the EC and *vice-versa*. Furthermore, it predicted denate cells in CA1, CA2, and entorhinal region.

On the other hand, SpatialPrompt achieves optimal predictions with an impressively shorter runtime of 2 minutes.

Similarly, spatial clustering was also performed for domain identification on both datasets. On the Visium cortex dataset, clusters labelled as 5, 2, 7, 4, and 14 were aligned sequentially, resembling the organisation of cortical layers L2/3 to L6 (**Fig. 3B-VIII**). Likewise, on the slide-seq dataset, precise clusters were obtained revealing the distinct anatomical features in the hippocampus (**Fig. 3D-VIII**). Specifically, cluster 10 predominantly consisted of CA1 and CA2 cell types, while cluster 7 majorly comprised CA3 cell types. Furthermore, cluster 2 was associated with DG cell-type and cluster 3 have EC cell-type.

### Application on spatial mouse kidney dataset using multiple scRNA-seq references

Spot deconvolution predicted by different methods varies significantly depending up on the choice of scRNA-seq reference^51,61^. To minimise this platform and batch effect arising due to different scRNA-seq references, SpatialPrompt employed several strategies in the feature selection, normalisation and spatial spot simulation steps as described in the method section. To assess the robustness of SpatialPrompt, we used the mouse kidney spatial dataset and two un-paired scRNA-seq reference datasets with accession id GSE157079 and GSE107585. The UMAP plot shows the distribution of different cell-types in the two scRNA-seq datasets (**Fig. 4A**).

**Fig. 4:**
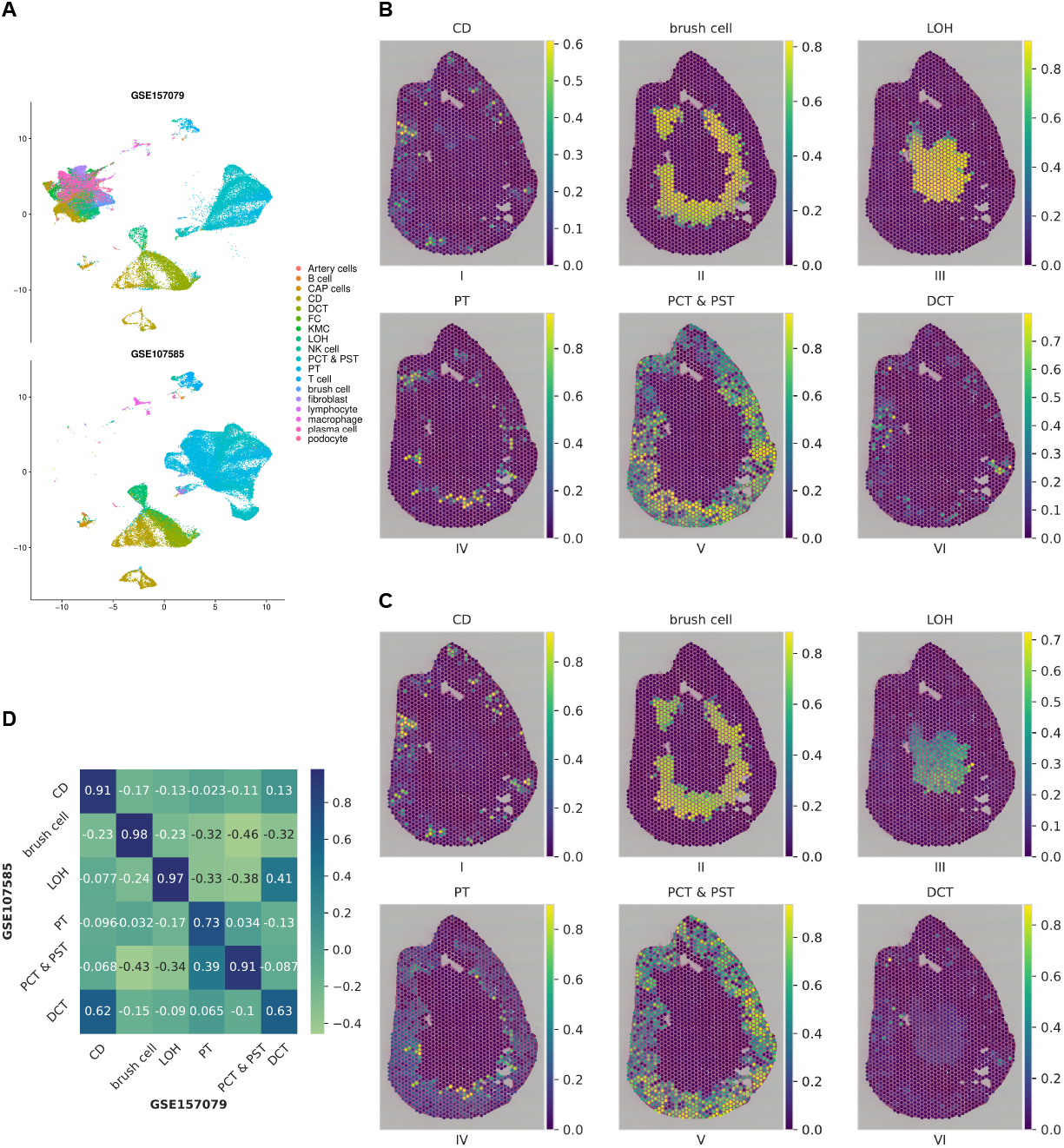
Consistent cell type inference prediction by SpatialPrompt by multiple references. (A) The UMAP plot of scRNA-seq references 1 and 2 having 18 cell types. (B and C) Spatial distribution of six major cell types predicted by SpatialPrompt using references 1 and 2. (D) Heatmap of the Pearson’s correlation coefficient obtained between the predictions obtained by using references 1 and 2.

Our analysis shows that SpatialPrompt was able to capture the major cell-types distribution in the kidney spatial dataset using two different references separately (**Fig. 4B & 4C**). The major cell types, including collecting ducts (CD), proximal convoluted tubule (PCT), and proximal straight tubule (PST) were observed in the outer cortex region (**Fig. 4B & 4C**). Additionally, brush cells and loop of Henle (LOH) cell-types extend from the outer cortex into the inner medulla region. **Fig. 4D** shows the heatmap of Pearson’s correlation coefficient (r^2^) calculated between the spot deconvolution predictions for 6 major cell-types obtained using reference 1 (GSE157079) and reference 2 (GSE107585), respectively. We observed r^2^ scores > 0.9 for 4 cell-types, namely, CD, brush cell, LOH, PCT, and PST cell-types. These results indicate an impressive robustness of SpatialPrompt while using multiple reference scRNA-seq datasets.

### Overview of SpatialPromptDB reference database

As SpatialPrompt tool can perform spot deconvolution using any scRNA-seq reference from the same tissue, we build SpatialPromptDB to provide compatible scRNA-seq references with cell-type annotations for seamless integration. Moreover, many of the scRNA-seq datasets available at the GEO database, the human cell atlas, and the Broad Institute single cell portal were found to have inconsistent data format. Most importantly, most of these databases either do not provide cell type annotations or accessing them is quite complicated. To overcome these issues, we manually curated 41 scRNA-seq datasets with their cell-type annotations from different tissues for mouse and human. All scRNA-seq datasets were pre-processed as described in the method section. In addition, the cell type annotations provided by the respective studies were manually verified using the CellMarker database^49^. All the reference datasets can be downloaded by the user either from the SpatialPromptDB database or by using our python package (https://github.com/swainasish/SpatialPrompt).

## Discussion

Advancements in spatial technologies during recent times have enabled researchers to sequence tissue samples while preserving their spatial coordinates. These spatial datasets have huge potential in tracking disease progression, studying developmental processes, exploring cellular crosstalk, and many more. However, commercially available sequencing-based spatial technologies such as ST and 10X Visium^3,5^ have low resolution of 10-100 *μm*, where each spatial spot consists of multiple cells. To understand the complex tissue structure, deconvolution of spatial spots is required to know the cell-type distribution. Furthermore, accurate spatial domain identification that reflects the biological reality remains a challenge in spatial transcriptomics.

Several computational methods have been developed for spot deconvolution and domain identification^11,14,16,22,23^ in spatial transcriptomics. However, most of the existing methods do not leverage the valuable spatial information. Also, recently developed domain identification methods that utilise spatial coordinates run extremely slow for larger datasets. To address these key issues, we introduced SpatialPrompt, a spatially aware, scalable and accurate cell-type deconvolution and domain identification method for spatial transcriptomics. SpatialPrompt uses a spatial simulator that simulate spatial spots using the reference scRNA-seq dataset and allow various criteria to match the characteristics of a real spatial data (**Fig. 1B)**. SpatialPrompt employs an iterative approach inspired from the message-passing layer in the GNN to incorporate the local microenvironment information into the spatial data. Moreover, a NRR model is used to learn the local microenvironment gene expression patterns from the real spatial data. SpatialPrompt uses KNN regression and k-means clustering for spot deconvolution and domain identification, respectively. The former learns the cell type specific gene signatures from the spatial embedded simulated matrix and predicts the cell type proportions in the real spatial data.

Our benchmarking analysis clearly showed that SpatialPrompt outperform the existing methods for both spot deconvolution and domain identification. We applied our proposed method on diverse datasets to explore its robustness. For initial benchmarking study, we used the human DLPFC Visium dataset containing the gold standard manual annotation of 7 cortical layers. Noteworthy, SpatialPrompt achieved the highest mean AUROC of 0.72 for predicting cortical layer specific 10 excitatory neuron cell-types. Among the 5 deconvolution methods benchmarked in our study, the Cell2location was the only method that come close to SpatialPrompt having mean AUROC of 0.60. Next, we applied SpatialPrompt and Cell2location to the mouse cortex and hippocampus datasets generated using low resolution Visium and high-resolution slide-seq techniques, respectively. In comparison to Cell2location, our tool successfully captured the cell type organisation from L2/3 to L6 layers in the mouse cortex (**Fig. 3B**). A previous study mentioned that most of the deconvolution tools applied to the same Visium dataset falsely predicted the L2/3 cell type in the centre of the cortex region which spans caudoputamen and nucleus accumbens area^57^. We noticed that the same false prediction made by the Cell2location tool in our analysis (**Fig. S1**). Conversely, SpatialPrompt predicted an almost negligible presence of L2/3 cells in the cortical centre region (**Fig. 3B-I**). Similarly, in the hippocampus dataset, SpatialPrompt accurately captured the anatomical features of the major cell-types. Furthermore, we benchmarked the domain identification performance of six different tools using the DLPFC dataset. Again, SpatialPrompt showed best performance with NMI score of 0.70 (**Fig. 2D**). We also noticed that BayesSpace and SEDR performed well with NMI score of 0.65, but their runtime was extremely high (**Table. 4**) and sensitive to hyperparameters.

The key feature of SpatialPrompt is its ability to scale for larger datasets. The runtime of all the chosen methods was compared on slide-seq dataset by varying numbers (i.e., 5,000 to 50,000) of spatial spots. For spot deconvolution using a small dataset with 10,000 spots, SpatialPrompt took 44 seconds, while other methods except Tangram took more than 35 minutes. For a large dataset with 50,000 spots, SpatialPrompt took only 2 minutes, while Tangram and Cell2location took 36 minutes and 4.4 hours, respectively. Similarly, for domain identification, SpatialPrompt was able to identify domains for 50,000 spots within 90 seconds. Overall, our tool was 44 to 150 times faster compared to other methods. These finding indicate that SpatialPrompt is more accurate and highly scalable compared to the existing methods.

A major problem in the spatial transcriptomic is the inconsistent spot deconvolution prediction up on using different scRNA-seq references. Therefore, we applied SpatialPrompt on a kidney Visium dataset using two different scRNA-seq references. Both chosen references were collected from two different platforms and locations. Interestingly, despite having technological and environmental differences in references, SpatialPrompt was able to capture the major cell-type distribution in the kidney spatial data for both scRNA-seq references (**Fig. 4**). This analysis shows the robustness of SpatialPrompt on multiple different scRNA-seq references. Thus, a researcher does not need to perform separate scRNA-seq experiment for the purpose of spot deconvolution while using SpatialPrompt. Moreover, we built SpatialPromptDB database that provides manually curated scRNA-seq reference datasets for seamless integration using our SpatialPrompt tool. Our database harbours more than 40 scRNA-seq datasets from humans and mice with manually curated cell-type annotations that could be directly used in the SpatialPrompt tool.

Further, in order to use SpatialPrompt for spot deconvolution, a few important considerations should be followed to optimise the performance and accuracy of SpatialPrompt. Firstly, the scRNA-seq reference dataset should capture a diversity of cell-types and expression profiles. Wherever feasible, the cell-type annotations provided by the reference database should be used for spot deconvolution. Secondly, the number of neighbours is a crucial hyperparameter should be carefully determined. For Visium and ST data analysis, a value in the range 20 to 30 is recommended for this hyperparameter, while for higher resolution data, a value from 50 to 60 is suggested.

In summary, our extensive evaluations on multiple datasets demonstrate the robustness and superior performance of SpatialPrompt in inferring the spatial distributions of cell-types and domains. Moreover, we have shown that SpatialPrompt exhibits remarkable scalability. It can perform spot deconvolution and domain identification merely within 120 seconds even for a large dataset with 50,000 spots. We built SpatialPromptDB, a public database, that hosts more than 40 scRNA-seq datasets from humans and mice with manually curated cell-type annotations which could be directly used with our tool. Based up on these outcomes, we anticipate that SpatialPrompt will emerge as a preferred method for the researchers targeting to perform spot deconvolution and domain detection in large-scale spatial datasets. Our tool is available as a Python package which can be easily integrated with Scanpy and other popular pipelines.

## Declaration

### Ethical Approval

Not applicable.

### Competing interests

The authors declare that there are no competing financial interests.

## Author Contributions

AKS performed data analyses; VP developed the database; JS contributed to the manuscript writing; PY designed and supervised the study; AKS, JS, and PY wrote the first manuscript draft; all authors contributed by comments and approved the final manuscript.

## Data and code availability

Nine publicly available datasets were utilised in this study. First, the spatial human DLPFC dataset was retrieved from the LIBD database (https://research.libd.org/spatialLIBD/), and the corresponding snRNA-seq was accessed from GEO (accession id: GSE144136). The second spatial mouse Visium cortex dataset retrieve from the 10X genomics database (https://www.10xgenomics.com/resources/datasets); corresponding scRNA-seq accessed from GEO (accession id: GSE71585) for our analysis, we used the raw processed data from https://satijalab.org/seurat/articles/spatial_vignette.html. The third spatial slide-seq dataset and the corresponding scRNA-seq retrieved from the Broad Institute single-cell portal (https://singlecell.broadinstitute.org/single_cell/study/SCP948/robust-decomposition-of-cell-type-mixtures-in-spatial-transcriptomics). The Last spatial mouse kidney data downloaded from STOmicsDB (https://db.cngb.org/stomics/datasets/STDS0000121) and two reference scRNA-seq data accessed from GEO (accession id: GSE157079, 107585).

SpatialPrompt Python package and benchmark scripts are available at: https://github.com/swainasish/SpatialPrompt. Detailed documentation, tutorials, and SpatialPromptDB database are available at: https://swainasish.github.io/SpatialPrompt/.

## Funding and acknowledgements

This project was supported by the seed grant (project number I/SEED/PY/20200037) funded by Indian Institute of Technology, Jodhpur and returning expert grant (S/GIZ/PY/20220140) funded by Deutsche Gesellschaft für Internationale Zusammenarbeit GmbH.

### URLs

CARD: https://yingma0107.github.io/CARD/documentation/04_CARD_Example.html. *Cell2location:* https://cell2location.readthedocs.io/en/latest/notebooks/cell2location_tutorial.html.

*Tangram*: https://github.com/broadinstitute/Tangram.

*SPOTlight*: https://marcelosua.github.io/SPOTlight/.

*RCTD:* https://github.com/dmcable/spacexr.

*STAGATE:* https://github.com/zhanglabtools/STAGATE.

SEDR: https://github.com/JinmiaoChenLab/SEDR.

SpaGCN: https://github.com/jianhuupenn/SpaGCN.

## Abbreviations

scRNA-seq: single-cell RNA sequencing
snRNA-seq: single-nucleus RNA sequencing
FISH: fluorescence in situ hybridisation
DLPFC: dorsolateral prefrontal cortex
NRR: non-negative ridge regression
GEO: gene expression omnibus
AUROC: area under the receiver operating characteristics
NMI: normalised mutual information
KNN: k-nearest neighbours
GNN: graph neural net
EC: entorhinal cortex

## Notes

### Competing Interest Statement

The authors have declared no competing interest.

