## Supplementary Data for "SpatialPrompt: spatially aware scalable and accurate tool for spot deconvolution and clustering in spatial transcriptomics"

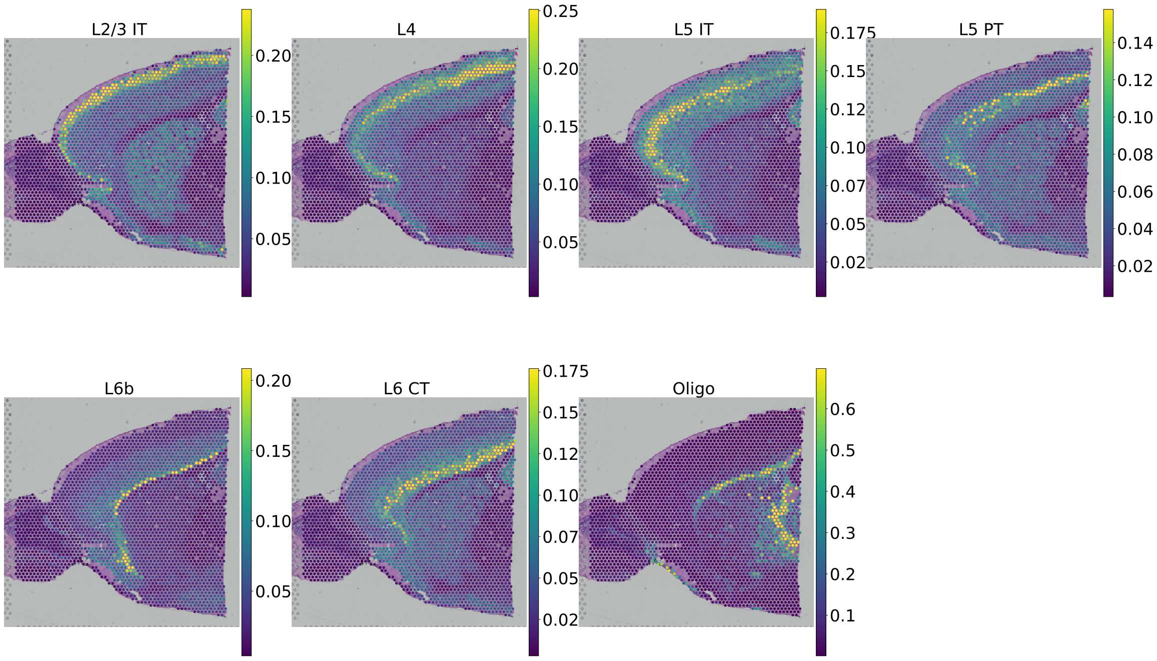


Fig. S 1: Spatial mapping of major cell types in mouse Visium cortex dataset by Cell2location tool.

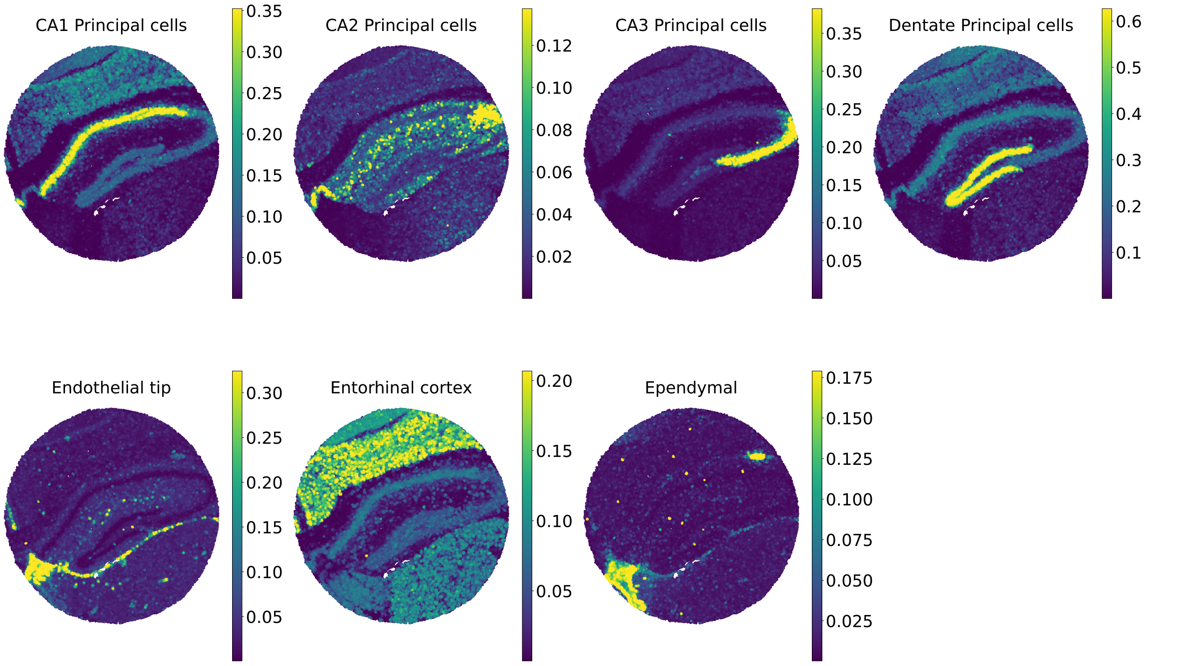


Fig. S 2: Spatial mapping of major cell types in mouse Slide-seq hippocampus dataset by Cell2location tool.

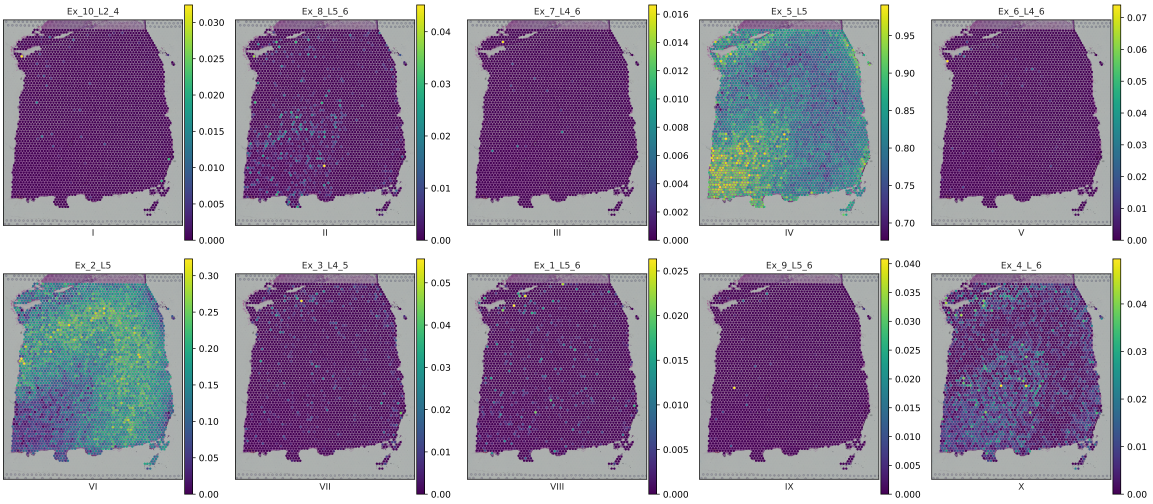


Fig. S 3: Spatial mapping of major cell types in human DLPFC dataset by CARD tool.

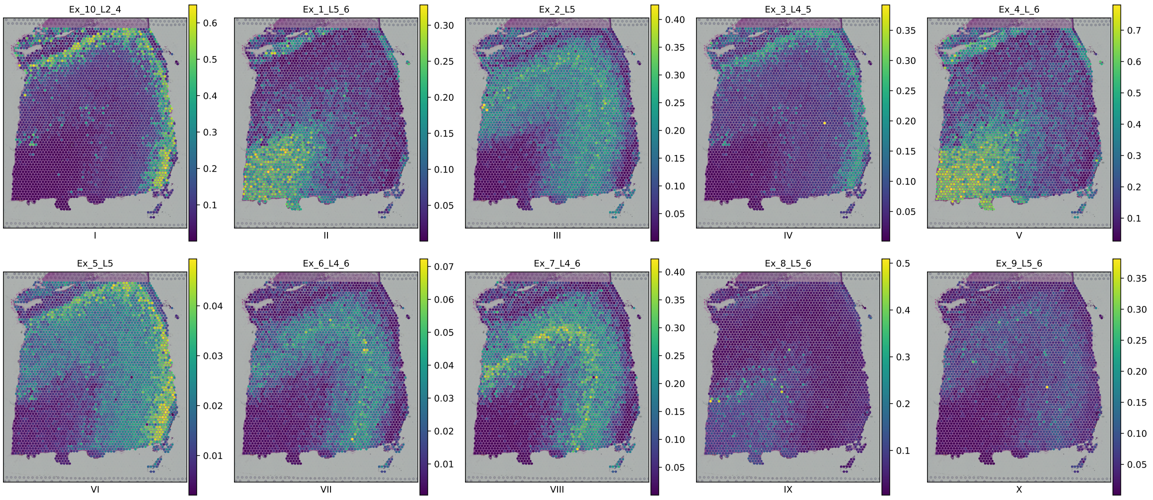


Fig. S 4: Spatial mapping of major cell types in human DLPFC dataset by Cell2location tool.

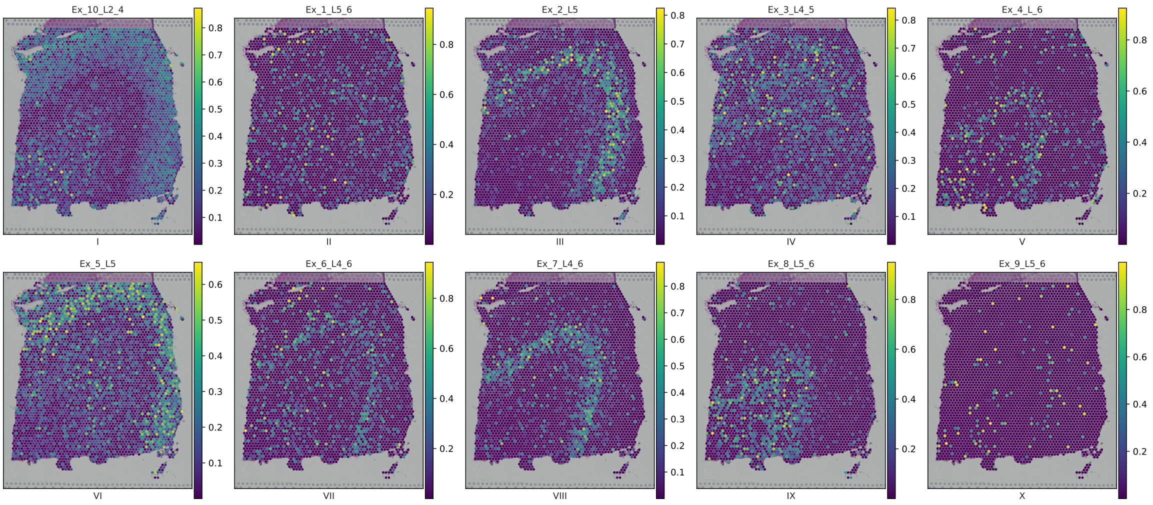


Fig. S 5: Spatial mapping of major cell types in human DLPFC dataset by Tangram tool.

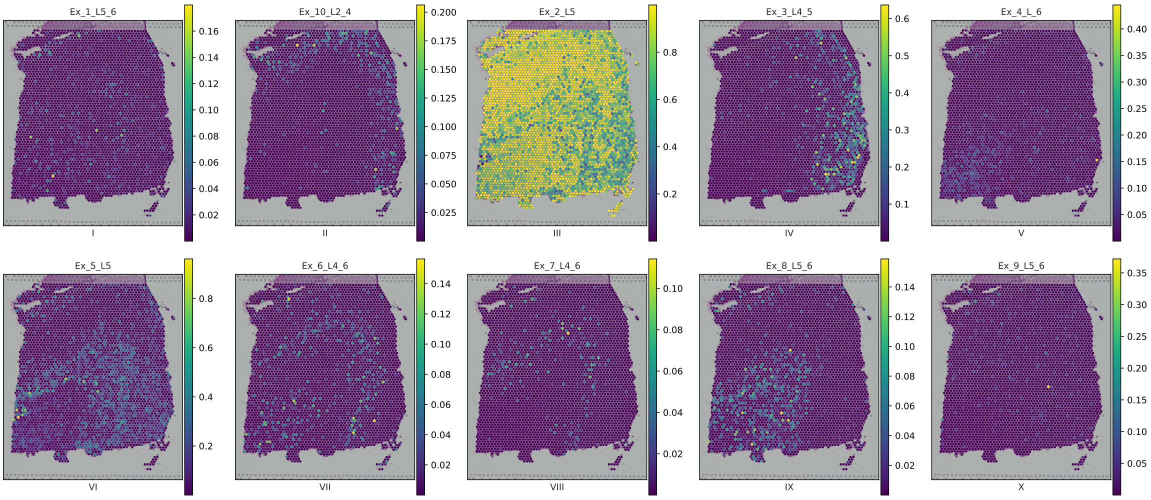


Fig. S 6: Spatial mapping of major cell types in human DLPFC dataset by RCTD tool.
